# The canonical Wnt signaling pathway is not a promising therapeutic target for PNS regeneration enhancement

**DOI:** 10.1101/2022.05.19.492703

**Authors:** Nikita Mehta, Maia Vardy, Benayahu Elbaz

## Abstract

PNS injury initiates transcriptional changes in Schwann cells, satellite glial cells and PNS neurons that facilitate regeneration. The signaling pathways that control these transcriptional changes are not fully understood. The canonical Wnt signaling pathway is active during early stages of PNS development, where it controls radial axonal sorting and the onset of PNS myelination. Upon PNS injury, the Wnt signaling pathway is re-activated, suggesting that Wnt signaling plays an important role in PNS regeneration. To explore the potential of the Wnt pathway as a therapeutic target for enhancement of PNS recovery, we used a combination of genetic and pharmacological approaches to either activate or inhibit the Wnt signaling pathway during PNS recovery. We found that manipulating the Wnt signaling pathway does not alter PNS regeneration. Our data suggests that the Wnt signaling pathway is not a strong therapeutic target for the enhancement of PNS regeneration.

## Introduction

During embryonic stages, immature Schwann cells, derived from Schwann cell precursor cells, deposit basal lamina around axon bundles, insert Schwann cell processes into the bundles and segregate large axons to the periphery, in a physiological process termed radial axonal sorting. Following this, Schwann cells myelinate large caliber axons in a 1:1 ratio, while small caliber axons remain unmyelinated in Remak bundles (Feltri et al., 2016). We and others have shown that the Wnt signaling pathway plays an important role in early PNS development by controlling radial axonal sorting and the onset of PNS myelination (Elbaz et al., 2016; Grigoryan et al., 2013).

In mammalians, axons and Schwann cells in the PNS are unique in their ability to recover from nerve injury. PNS injury initiates transcriptional changes in both myelinating and non-myelinating Schwann cells, resulting in a new transcriptional state that promotes regeneration. Schwann cells that lose their axon-glia interactions upon injury transdifferentiate and transform to repair Schwann cells that can aid in axonal regeneration (Jessen & Mirsky, 2016, 2019). This transcriptional reprograming is controlled mainly by the early transcription factor c-Jun (Arthur-Farraj et al., 2012), the transcription factor STAT3 (Benito et al., 2017), by chromatin modifications (Ma & Svaren, 2018), and by Merlin (Mindos et al., 2017). Upon completion of neuronal regeneration and reinnervation, the original Schwann cell identity is reestablished, and Schwann cell axon-glia interactions are restored (Jessen & Mirsky, 2016, 2019). In parallel, PNS neurons responds to injury by a transcriptional reprograming that suppresses their sensory neuron cell identity and promotes neuronal regeneration. In sensory neurons, this reprograming depends on the early transcription factor ATF3 (Renthal et al., 2020).

Transcripts that constitute the canonical Wnt signaling pathway are induced following nerve injury (Duraikannu et al., 2018; Vliet et al., 2021; Wang et al., 2016; Yao et al., 2012), suggesting that Wnt signaling may regulate the transcriptional changes that modulate PNS regeneration. Wnt signaling pathway activity is mediated by β-catenin, which is constitutively expressed. Upon the interaction between the Wnt ligand, Frizzled receptor, and either the LRP5 or LRP6 receptor, the Wnt signaling pathway is activated, and β-catenin mediates gene expression through its binding to the transcription factors LEF1, TCF1, TCF3, and TCF4 (Kim et al., 2013; Näthke, 2006). In the absence of the Wnt ligand, β-catenin is targeted for degradation by the β-catenin destruction complex, which is comprised of four proteins: Glycogen Synthase Kinase 3 (GSK-3β), Casein Kinase 1 (CK1), Axin2 and Adenomatous Polyposis Coli (APC). In the presence of the Wnt ligand, the β-catenin destruction complex disassembles, allowing for accumulation and translocation of β-catenin to the nucleus (Kim et al., 2013; Näthke, 2006).

During developmental PNS myelination, Wnt signaling is activated in a temporal manner. Wnt activity in the sciatic nerve of mice begins at embryonic day 12.5, reaches a peak between embryonic day 15.5 and 17.5, and starts declining at postnatal day 15, once developmental myelination has been completed (Grigoryan et al., 2013). Developmental PNS myelination is affected by both genetic and pharmacological inhibitors and activators of the Wnt signaling pathway (Elbaz et al., 2016; Grigoryan et al., 2013). APC ablation during developmental PNS myelination results in the constitutive activation of the Wnt signaling pathway and mild hypomyelination characterized by reduced myelin thickness and shorter internodes, which disrupts motor function of APC-ablated mice (Elbaz et al., 2016).

Based on the induced expression of transcripts of the Wnt signaling pathway following PNS injury (Duraikannu et al., 2018; Vliet et al., 2021; Wang et al., 2016; Yao et al., 2012), we sought to explore the effects of Wnt signaling pathway manipulation on the course of PNS recovery. To activate Wnt signaling, we used the Cre-Lox system to ablate *APC* specifically from Schwann cells. To inhibit the Wnt signaling pathway, we used the porcupine inhibitor LGK974 and the drug Pyrvinium (Liu et al., 2013; Thorne et al., 2010), both of which are well characterized Wnt signaling inhibitors. Our genetic and pharmacological studies suggest that manipulating the Wnt signaling pathway does not change PNS regeneration.

In humans, PNS regeneration is slow and inefficient, and with age, the ability of the PNS to recover from injury declines with alarming celerity (Scheib & Höke, 2013). Therefore, a comprehensive understanding of the molecular control of PNS regeneration is crucial for our ongoing efforts to expedite PNS recovery. Our results suggest that the Wnt signaling pathway is not a promising therapeutic target for the enhancement of PNS regeneration.

## Materials and Methods

### Animals

Mice were housed and studied according to the institutional Animal Care and Use Committee (IACUC) guidelines. Schwann cell-specific conditional knockout *APC* mice (*APC^lox/lox^;PLP-CreERt*) were generated by mating mice carrying an *APC* allele in which exon 14 is flanked by *loxP* sites (Shibata et al., 1997) with the *PLP-CreER^t^* transgenic mouse line (Doerflinger et al., 2003). Littermate *APC^lox/lox^* mice served as controls. Mice were genotyped by PCR analysis of tail genomic DNA using the primers P3 (5’-GTTCTGTATCATGGAAAGATAGGTGGTC-3’) and P4 (5’-CACTCAAAACGCTTTTGAGGGTTGATTC-3’). The excised allele was detected using primers P5 (5-GAGTACGGGGTCTCTGTCTCAGTGAA-3’) and P3 (Shibata et al., 1997).

The *PLP-CreER^t^* reporter line was generated by mating *ROSA26-stop-EYFP* mice (Srinivas et al., 2001) with the *PLP-CreER^t^-Cre* transgenic mouse line (Doerflinger et al., 2003). The TCF/Lef:H2B-GFP reporter mice (Ferrer-Vaquer et al., 2010) were purchased from Jackson Labs (Stock #: 013752).

### Nerve Crush Surgery

We performed a nerve crush surgery on 8–10-week-old *PLP-CreER^t^-APC*^Lox/Lox^ mice and Cre negative *APC*^Lox/Lox^ mice (used a control) and on the TCF/Lef:H2B-GFP reporter mice as previously described (Goodrum et al., 1995; Rath et al., 1995; Popko et al., 1993). Mice from the inducible Cre line were injected 4-hydroxytamoxifen for five consecutive days two weeks prior to the nerve crush surgery as detailed below. To perform the surgery, the right sciatic nerve was exposed at its exit from the vertebral canal. Jeweler’s forceps were used to crush the nerve at the foramen cross site. The crush was maintained for 15 seconds, such that nerve injury was sustained, while the epineurium remained intact (Sunderland, 1990). As a control, a sham procedure (skin incision) was administered to the contralateral leg. The injured limb lost function immediately, which was generally regained by 4 weeks post-crush injury. This pattern corresponded to normal nerve repair in adult mice. We engraved hemostatic forceps with a mark 1 millimeter from the tip to ensure that each crush was reproducible. The outermost point at the site of crush was placed in line with the 1-mm mark, which ensured uniform width during the crush.

### Drug Treatment

To ablate APC, 6-8 weeks-old mice were intraperitoneally injected with 0.1mL of 10mg/mL 4-hydroxytamoxifen (Sigma Aldrich) dissolved in a solution of DMSO, Ethanol, and Sunflower oil for five consecutive days. Animals were treated immediately following nerve crush daily with either 5mg/kg of Pyrvinium (Med Chem Express, HY-A0293) for seven days or treated with 3mg/kg of LGK974 (Med Chem Express, Cat: HY-17545) through oral gavage for a period of seven days post-crush (Liu et al., 2013). The drug regime for the LGK974 and Pyrvinium was based on recent studies who demonstrated activation of the Wnt signaling key ligands Wnt5b and Wnt4 at about 3 days post injury (Duraikannu et al., 2018; Vliet et al., 2021).

### EM Imaging and Myelin Morphometry

Animals were anesthetized with avertin (0.5 mg/g) through intraperitoneal injection and intracardially perfused (through the ventricle) with saline for 10 minutes, followed by a perfusion with EM buffer (2.5% glutaraldehyde and 4% paraformaldehyde in a 0.1M sodium cacodylate buffer). Sciatic nerves were harvested and post-fixed in EM buffer and embedded in an epoxy resin. Thin cross-sections of the sciatic nerve were cut, double stained with uranyl acetate and lead citrate, and observed on a Philips CM 10 electron microscope. G-ratios were calculated from the sciatic nerves of at least three mice per genotype. The g-ratios were calculated as axon diameter divided by fiber diameter as described previously (Auer, 1994).

### Immunohistochemistry

Animals were anesthetized with avertin (0.5 mg/g) through intraperitoneal injection and intracardially perfused (through the ventricle) with saline for 10 minutes, followed by a perfusion with 4% paraformaldehyde for 10 minutes. Sciatic nerves were harvested and post-fixed in 4% paraformaldehyde for 2 hours and left in a sucrose and 0.1% sodium azide solution overnight. Cross-sections of tissue were cut (10um thickness) and immunostained. The antibodies used were against GFP (Invitrogen, A21311, 1:500), p75NTR (Cell Signaling Technology, 8238T, 1:250) and MBP (BioRad, aa82-87 1:500). Confocal images of the sciatic nerves were taken with a Leica SP2-AOBS laser scanning microscope using a 20x, 40x or 60x objective.

### Behavioral analyses

Gait analysis was performed using the DigiGait (Mouse Specifics Inc., Boston, MA) imaging system. In this system, a high-speed digital camera captured the movement of the paws as the mouse walked on a treadmill belt, and simultaneously analyzed various gait parameters including Sciatic Functional Index (SFI). Each animal was tested on the DigiGait treadmill ten times on the day of testing, with 2-minute breaks in between. Once the animals walked consistently for at least 3 complete strides, data were collected.

Motor coordination and balance were examined using the accelerating rotarod (Colombus Instruments). Rotarod analysis was performed essentially as described in (Traka, 2019; Traka et al., 2010). *PLP-CreER^t^-APC^Lox/Lox^* mice and Cre negative *APC^Lox/Lox^* mice (used as control) were trained to move on the rotarod at a constant speed (5□rpm), and then were tested once a week (4 trial sessions) for 7 consecutive weeks. The time taken to fall off the rod in accelerating speed mode from 5-65 rpm (latency) was recorded during 5-minute trial sessions.

### Nerve Electrophysiology

To examine peripheral nerve function, animals were subjected to electrophysiological examination. A Nicolet Viking Quest Laptop System (VikQuest Port 4ch-7) was used for these studies. Recording needle electrodes were placed subcutaneously in the footpad. Supramaximal stimulation of the sciatic nerve with a 0.1–0.2 millisecond pulse, stimulating distally at the ankles and proximally at the sciatic notch with electrodes. Latencies, conduction velocities, and amplitudes of compound muscle action potentials were measured to identify axonal and myelin abnormalities.

### Data Analysis

Either two-tailed unpaired student’s t-tests or two-way ANOVA for multiple comparisons was used for statistical analysis of the data.

## Results

### APC is not important for PNS myelin maintenance

APC is constitutively expressed in the PNS, by both Schwann cells and DRG neurons (Duraikannu et al., 2018). We therefore sought to assess its importance in PNS myelin maintenance before we determined its importance in PNS injury. We have shown previously that Cre-mediated recombination of APC using the constitutive Schwann-cell specific PO-Cre line (Feltri et al., 1999) results in developmental loss of APC in Schwann cells and canonical Wnt/β-catenin signaling pathway activation (Elbaz et al., 2016). To assess the importance of APC in PNS myelin maintenance in adults while leaving the developmental PNS myelination intact, we used the inducible *PLP-CreER^t^* line, which drives recombination of genes expressed under the PLP promoter, which is specific to oligodendrocytes in the CNS and Schwann cells in the PNS (Doerflinger et al., 2003). This line was bred to the APC line carrying an *APC*^Lox/Lox^ allele in which exon 14 was flanked by loxP sites (Shibata et al., 1997), resulting in the generation of a *PLP-CreER^t^* - *APC*^Lox/Lox^. Upon tamoxifen-induced *PLP-CreER^t^* mediated recombination, axon 14 of the *APC* gene is excised, preventing APC expression in Schwann cells.

To confirm tamoxifen-induced Cre-mediated recombination in Schwann cells in the sciatic nerve of the *PLP-CreER^t^* mice, we bred the *PLP-CreER^t^* line with the reporter mouse line ROSA26-stop-EYFP (Srinivas et al., 2001), in which the expression of EYFP is prevented by upstream “lox-stop-lox” (LSL) cassette. Upon tamoxifen-induced *PLP-CreER^t^* mediated recombination, the stop sequence is removed, and the expression of EYFP is induced. Tamoxifen-induced, *PLP-CreER^t^*-mediated recombination results in EYFP expression in both myelinating and non-myelinating Schwann cells in the *PLP-CreER^t^*-ROSA26-stop-EYFP line, demonstrating successful Cre mediated recombination in Schwann cells (Figure 1A).

**Figure 1:**
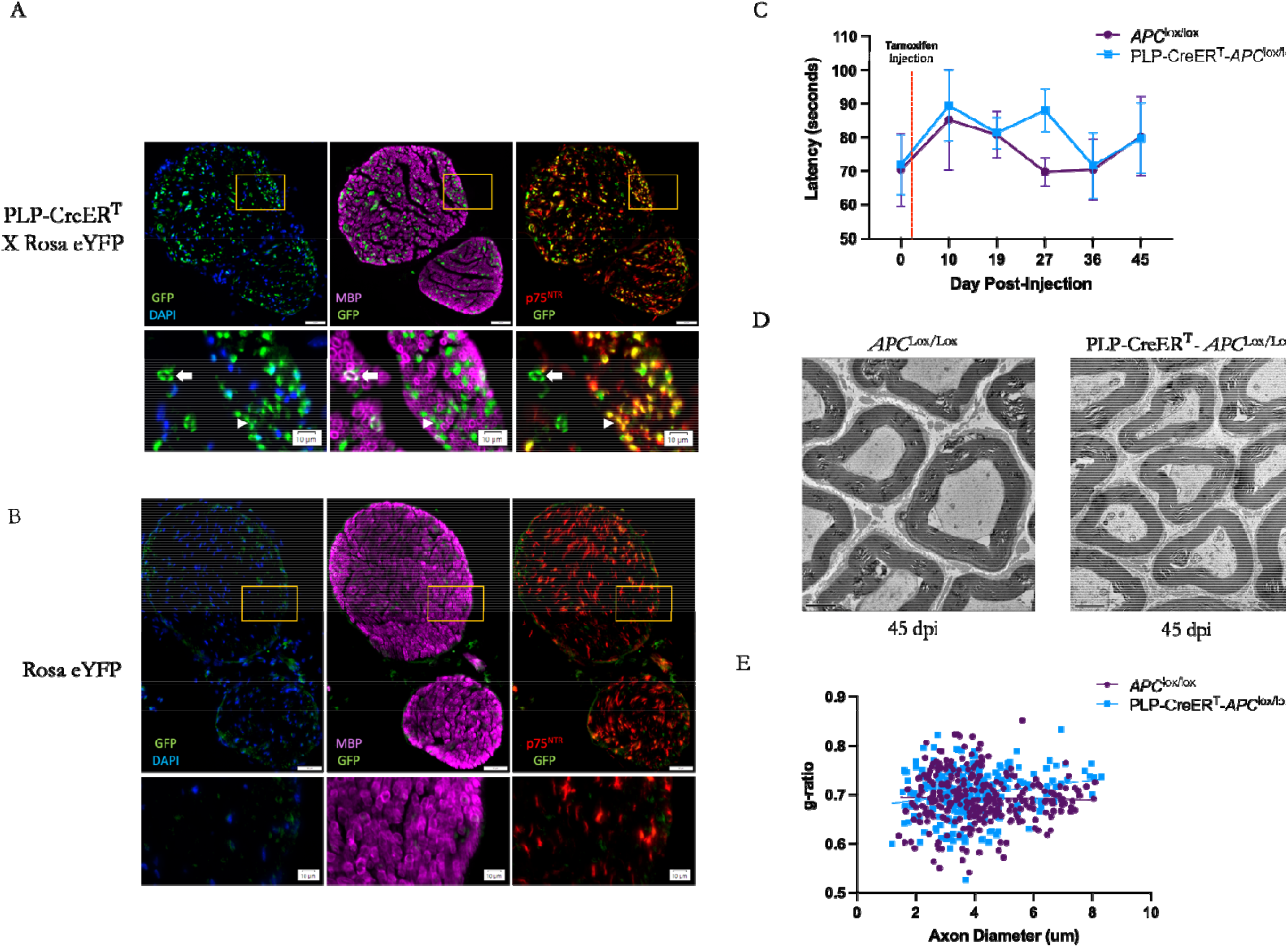
APC is not important for PNS myelin maintenance. **(A)** Eight-week-old *PLP-CreER^t^*; ROSA26-EYFP and Cre-negative ROSA26-EYFP mice (used as control) were injected with tamoxifen. Two weeks following injection, their sciatic nerves were harvested, and EYFP expression was detected using an anti-GFP antibody (green). Cre mediated EYFP expression was detected in in non-myelinating Schwann cells, labeled by p75^NTR^ (red) and in myelinating Schwann cells, labeled by MBP (pink). Lowe panel-higher magnification. White arrowhead mark EYFP positive cells that are positive for p75^NTR^, and white arrows mark EYFP positive cells that are positive for MBP. **(B)** Cre mediated EYFP expression was not detected in Cre-negative ROSA26-EYFP mice, used as control, that were injected with tamoxifen. **(C)** Eight-week-old *PLP-CreER^t^* - *APC*^Lox/Lox^ and Cre-negative *APC*^Lox/Lox^ mice (used as control) were injected with tamoxifen, following which an accelerating Rotarod assay was used to measure latency to fall. The mean Rotarod latency was 76.17 seconds for the Cre-negative *APC*^Lox/Lox^ mice and 80.37 seconds for *PLP-CreER^t^* - *APC*^Lox/Lox^ mice. The difference in the mean latencies of these two groups is not significant, with a p-value<0.3379. (D) Eight-week-old *PLP-CreER^t^* - *APC*^Lox/Lox^ and Cre-negative *APC*^Lox/Lox^ mice (control) were injected with tamoxifen. Forty-five days post injection (dpi) the sciatic nerves from these mice were harvested for EM analysis. Representative EM images of myelinated cells from sciatic nerve sections at 45 dpi are shown here. **(E)** Myelin thickness (g-ratio) was 0.70 in the *PLP-CreER^t^* - *APC*^Lox/Lox^ mice and 0.6934 in the Cre-negative *APC*^Lox/Lox^ mice. There is no significant difference in the g-ratio between these two groups, with a p-value<0.0613. Two-tailed unpaired t-tests were used to assess significance for Rotarod data and g-ratio data. Significance thresholds (not shown in figure): P > 0.05 (n.s.), P ≤ 0.05 (*), P≤ 0.01 (**), P≤ 0.001(***). For rotarod analysis, N = 12 mice; 7 *PLP-CreER^t^* - *APC*^Lox/Lox^ and 5 Cre-negative *APC*^Lox/Lox^. For g-ratio analysis, N = 6 mice; 3 mice/group; 291 Cre-negative *APC*^Lox/Lox^ axons and 200 *PLP-CreER^t^* - *APC*^Lox/Lox^ axons were analyzed.

To test the effect of Schwann cell-specific ablation of *APC* in adult *PLP-CreER^t^* - *APC*^Lox/Lox^ on PNS myelination and motor performance, eight-week-old *PLP-CreER^t^* - *APC*^Lox/Lox^ and Cre-negative *APC*^Lox/Lox^ mice (used as control) were injected with tamoxifen, following which an accelerating Rotarod assay was used to measure latency to fall over 45 days (Figure 1C). Over this period, the mean Rotarod latency was 76.17 seconds for the Cre-negative *APC*^Lox/Lox^ mice and 80.37 seconds for *PLP-CreER^t^* - *APC*^Lox/Lox^ mice. The difference in the mean latencies of these two groups was not significant, with a p-value<0.3379. At 45 days post-injection (dpi) the sciatic nerves from these mice were harvested for EM analysis (Figure 1D). EM images of myelinated cells from sciatic nerve sections and g-ratio calculations demonstrate that the myelin thickness (as measured by g-ratio), is not significantly different between the *PLP-CreER^t^* - *APC*^Lox/Lox^ (mean g-ratio = 0.70) and Cre-negative *APC*^Lox/Lox^ mice (mean g-ratio = 0.6934), with a p-value<0.0613. (Figure 1D-E). The results suggest that tamoxifen-mediated, Schwann cellspecific ablation of *APC* in adult *PLP-CreER^t^* - *APC*^Lox/Lox^ does not change the motor performance of the mice on the Rotarod (Figure 1B) and does not change either myelin integrity (Figure 1C) or thickness (Figure 1D-E). Taken together, this data suggests that *Apc* ablation in Schwann cells does not affect PNS myelin maintenance.

### Schwann cell-specific ablation of APC does not affect PNS regeneration following sciatic nerve crush

After establishing that Wnt signaling is likely not involved in PNS myelin maintenance, we turned our attention towards a potential role for Wnt signaling in regeneration. We used a combination of genetic and pharmacological approaches to study the effects of activating or inhibiting Wnt signaling during PNS regeneration following sciatic nerve crush. In line with the known role of APC as a Wnt signaling pathway inhibitor, we have shown previously that constitutive ablation of *Apc* in Schwann cells results in activation of the Wnt signaling pathway (Elbaz et al., 2016). Following from this finding, we activated the Wnt signaling pathway during PNS regeneration by ablating *Apc* from Schwann cells two weeks prior to sciatic nerve crush via tamoxifen injection. The experimental design of these studies is presented in Figure 2A. In these experiments, we ablated *Apc* from Schwann cells two weeks prior to sciatic nerve crush and analyzed myelin and functional regeneration at several time points during the recovery process. We found that nerve crush results in damaged myelin sheath and demyelinated axons at 14 days post-crush (Figure 2B). The sciatic nerve was largely remyelinated 28 days post crush (Figure 2B), although the myelin was thinner than the myelin in non-crushed legs. Importantly, *Apc* ablation did not result in aberrant remyelination and did not affect myelin thickness during the recovery (Figure 2C).

**Figure 2:**
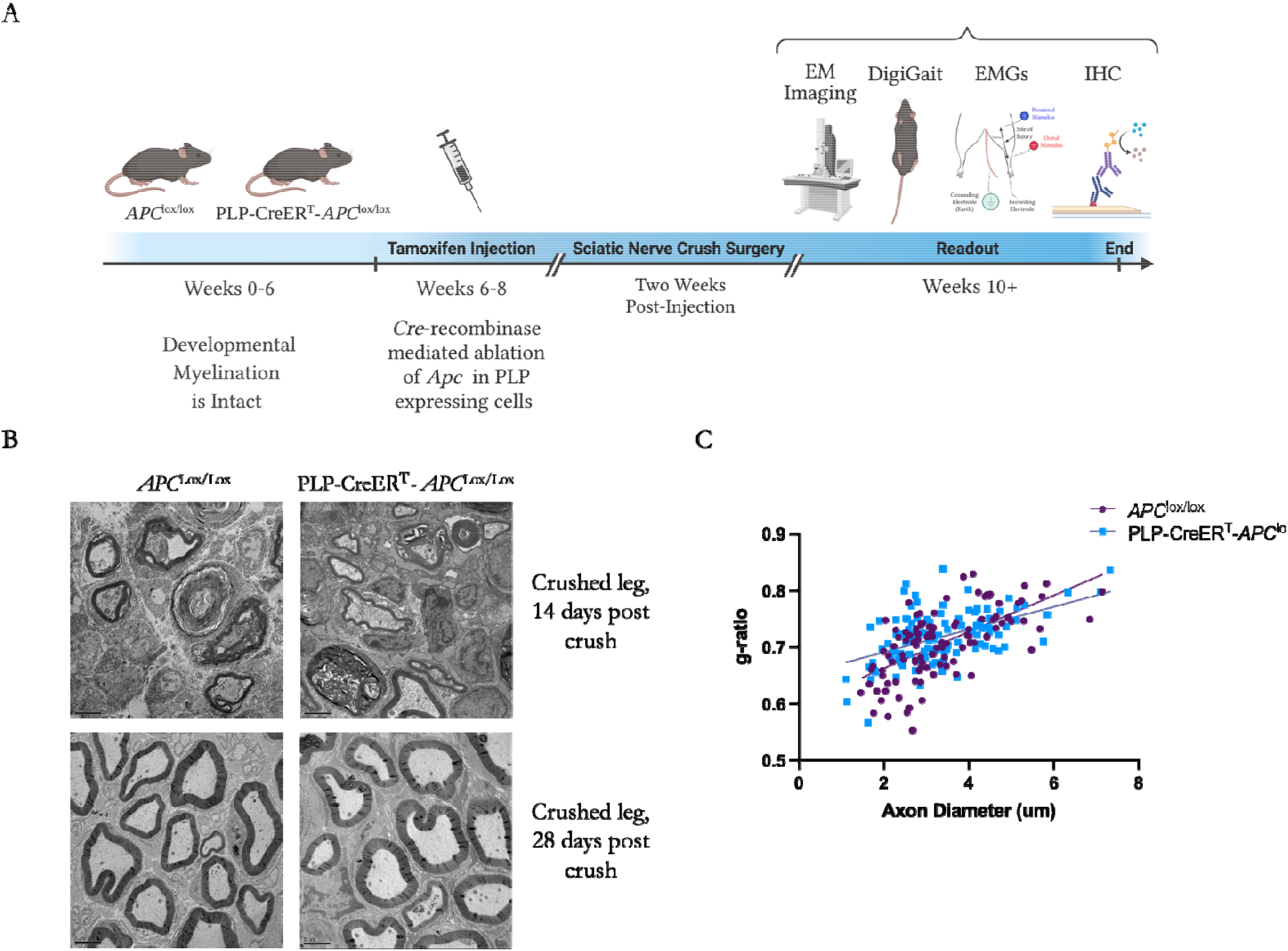
Schwann-cell specific ablation of APC does not affect myelin regeneration following sciatic nerve crush injury. **(A)** Schematic diagram of the experimental design. *PLP-CreER^t^* - *APC*^Lox/Lox^and Cre-negative *APC*^Lox/Lox^ (used as control) were injected with tamoxifen. Two weeks post injection, the sciatic nerve in the right limb was subjected to nerve crush injury (see Materials and Methods). Following crush injury, a series of experiments were performed to assess recovery, myelin regeneration, and electrophysiological properties of the regenerating nerve. (B) The sciatic nerves from a subset of these mice were harvest at 14- and 28-days post crush for EM imaging and analysis. Representative EM micrographs for *PLP-CreER^t^* - *APC*^Lox/Lox^ and Cre-negative *APC*^Lox/Lox^ mice are shown here. In both genotypes demyelination was visible at 14 dpi, and all the large caliber axons were remyelinated at 28 dpi. (C) Myelin thickness (g-ratio) as a factor of axon diameter was assessed based on the EM micrographs. There was no significant difference in mean g-ratio of *PLP-CreER^t^* - *APC*^Lox/Lox^ mice (0.7189) and Cre-negative *APC*^Lox/Lox^ mice (0.7068) at 28 days post-crush (two-tailed unpaired t-test, p<0.0935). N = Sections were taken from 8 mice, 4 mice/group; 224 axons distal to the site of injury were analyzed for both groups combined.

Next, functional recovery of the mice was measured using the DigiGait system (Figure 3). Mice exhibited characteristic hind limb clenching and paw abnormalities post-crush, reflecting the crush injury (Figure 3A-3F). While nerve crush injury reduced the paw width and paw length, which are measures of impaired gait (Figure 3H-I), there were no significant differences between the two groups in paw width or paw length at any time point (Figure 3G-H). Ablation of *Apc* did not change the functional recovery of the sciatic nerve, as measured by the Sciatic nerve Functional Index (SFI) (Figure 3I).

**Figure 3:**
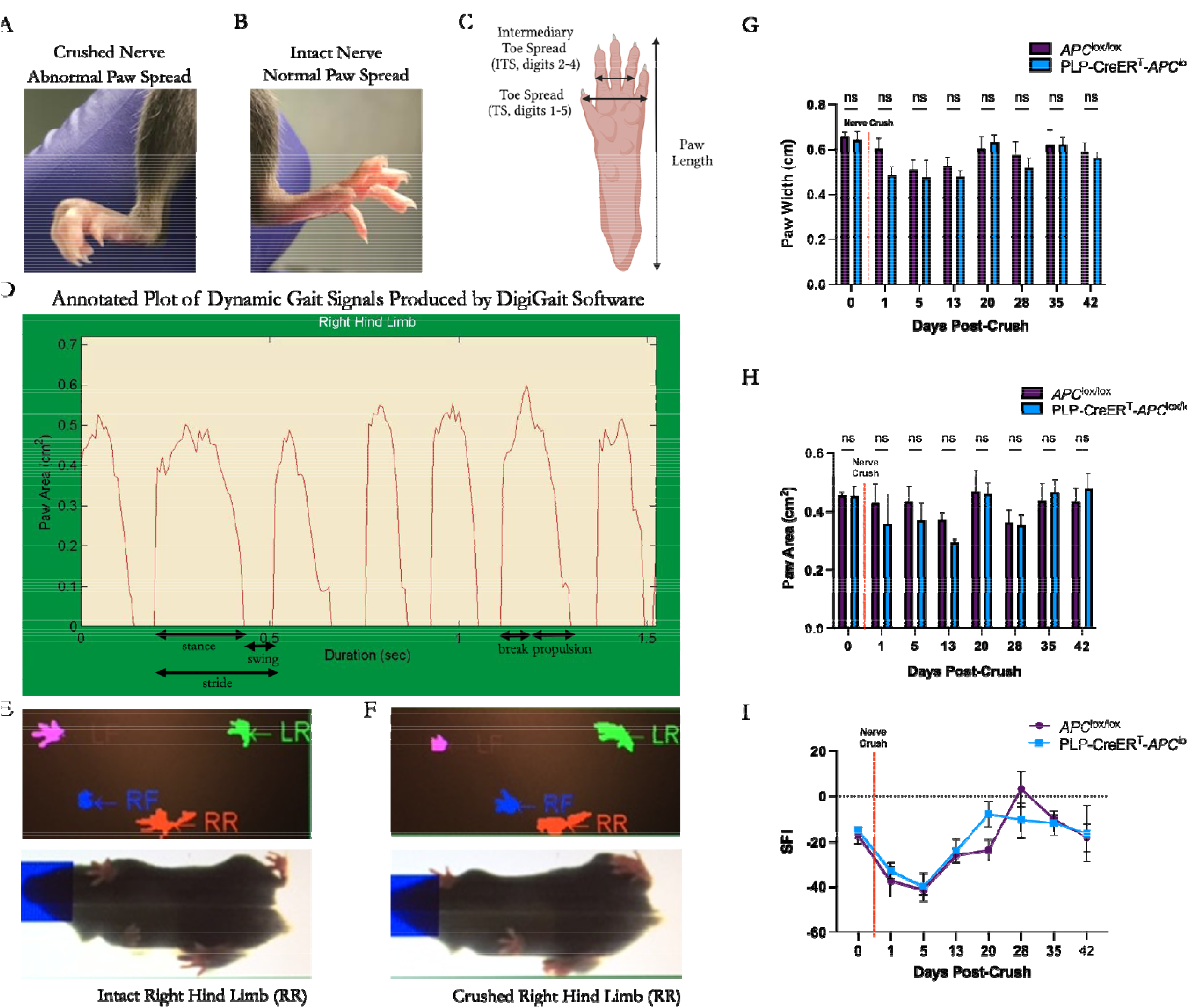
Schwann cell-specific ablation of APC does not change the functional recovery of the sciatic nerve following sciatic nerve crush injury. Eight-week-old *PLP-CreER^t^* - *APC*^Lox/Lox^ and Cre-negative *APC*^Lox/Lox^ mice (used as control) were injected with tamoxifen. Two weeks post injection, the sciatic nerve in the right limb was subjected to nerve crush injury (see Materials and Methods) and motor functions were studied. (A) Sciatic nerve crush resulted in abnormal paw spreading. **(B)** Normal paw spreading in non-injured leg. **(C)** Paw length, intermediary toe spread (ITS, digits 2-4), and toe spread (TS) were used to assess recovery from sciatic nerve crush injury. **(D)** DigiGait was used to assess motor function recovery for a 42-day post-crush injury period. An example of DigiGait-generated plot of dynamic gait signals (measured by paw area) over time. Stance, stride, swing, break, and propulsion are annotated on the plot. **(E-F)** Example of signals captured by DigiGait for mouse with crushed nerve in the right hind limb **(E)** and mouse with intact nerve in the same limb **(F). (G)** Paw width and **(H)** paw area was compared between the two groups. While both paw width and paw area were reduced upon sciatic nerve crush, no difference between the genotypes was observed. **(I)** Sciatic Nerve Functional Index (SFI) at different time points following injury. Sciatic nerve injury reduced the SFI, which is a geometric representation of the injured paw compared to the contralateral paw, however, there was no statistical difference between the two genotypes. Twoway ANOVA for multiple comparisons was used to assess significance. N = 4 mice per genotype.

Upon sciatic nerve injury, Schwann cells distal to the injury site undergo transcriptional reprograming and turn into repair Schwann cells. This process is accompanied by distal myelin loss, axonal loss and denervation. While measuring the compound muscle axon potential (CMAP), the loss of myelin is reflected by reduced nerve conduction velocity, and the axonal loss is reflected by reduced amplitude. The electrophysiological properties of the regenerating nerve were not significantly altered when *Apc* was ablated (Figure 4). By 45 days post-crush, both the control and APC-ablated mice recovered such that pre-crush nerve properties (CV, distal amplitude) were not significantly different than 45 days post-crush (Figure 4C-D, 4F-G). Taken together, our electrophysiological and functional studies suggest that Schwann cell-specific ablation of APC does not change PNS regeneration.

**Figure 4:**
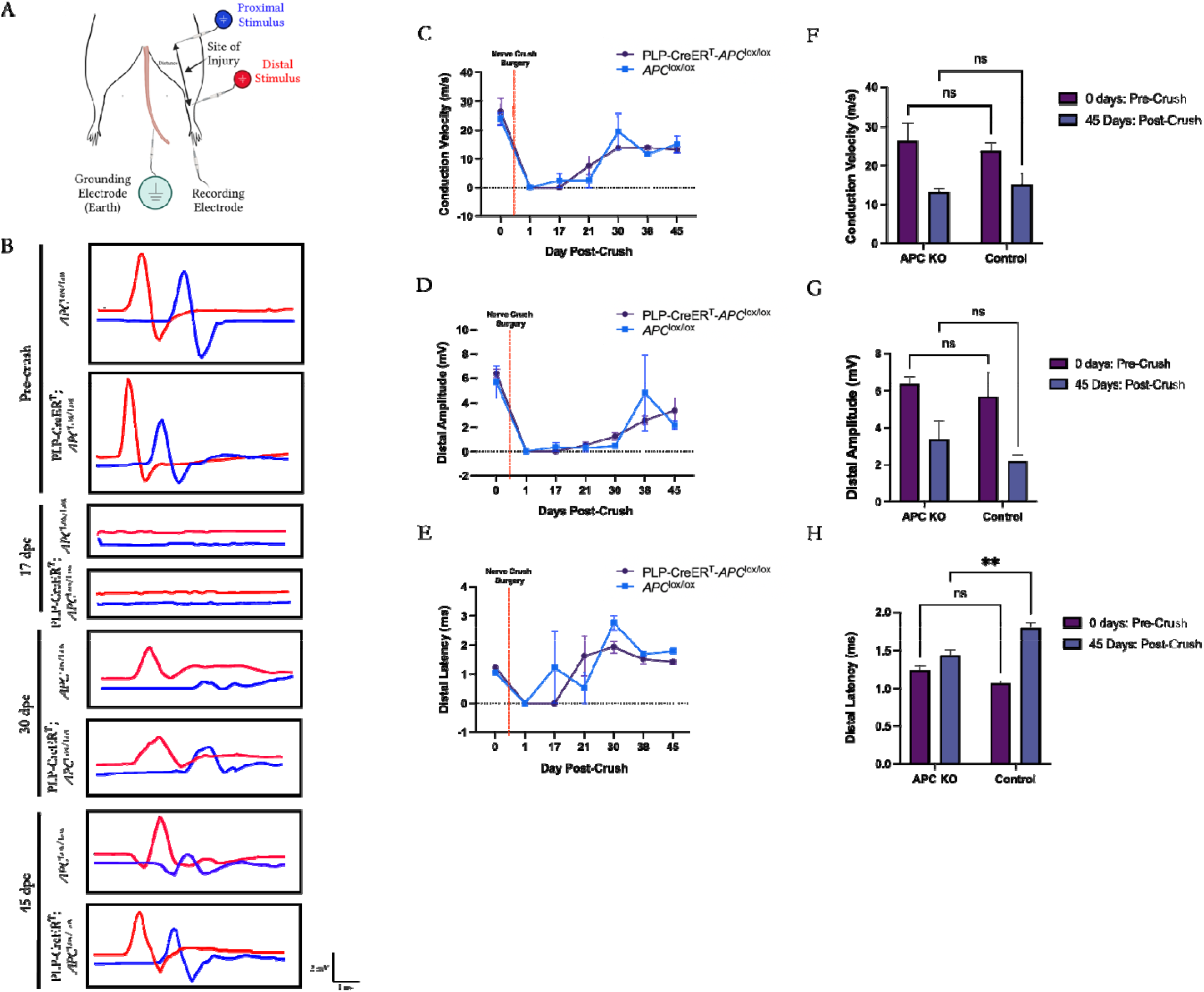
Schwann cell-specific ablation of APC does not alter the electrophysiological properties of the sciatic nerve following sciatic nerve crush injury. Eight-week-old *PLP-CreER^t^* - *APC*^Lox/Lox^ and Cre-negativc *APC*^Lox/Lox^ mice (used as control) were injected with tamoxifen. Two weeks post injection, the sciatic nerve in the right limb was subjected to nerve crush injury (see Materials and Methods). Following crush injury, the electrophysiological properties of the sciatic nerve were examined at different time points. (A) Diagram illustrating the recording setup used to measure the compound motor action potentials (CMAPs) in the sciatic nerve. **(B)** Representative traces of electrophysiological findings in control and *PLP-CreER^t^* - *APC*^Lox/Lox^ mice pre-crush, 17, 30 and 45 days post-crush injury. Distal CMAP trace is in red, proximal CMAP trace is in blue. **(C-E)** In both groups of mice, the conduction velocity (m/s), distal amplitude (mV), and distal latency (ms) could not be measured in sciatic nerves following crush injury, but by 45 days post-injury, both groups of mice demonstrated partial nerve recovery. No significant difference between the two genotypes was observed. **(F-H)** Quantified results. The conduction velocity and distal amplitude in *PLP-CreER^t^* - *APC*^Lox/Lox^ mice at 45 days was not significantly different when compared to Cre-negative *APC*^ox/Lox^ mice (control). The distal latency had significant but small reduction in the experimental group, which by itself is not indicative of recovery differences between the two groups. Two-way ANOVA for multiple comparisons was used to assess significance. Significance levels: P > 0.05 (n.s.), P ≤ 0.05 (*), P≤ 0.01 (**), N = 8 mice, 3 controls, and 5 *PLP-CreEP^t^* - *APC*^Lox/Lox^ mice.

### Pharmacological inhibition of the Wnt Signaling Pathway does not affect sciatic nerve regeneration

To inhibit the Wnt signaling pathway, we used the drugs LGK974 and Pyrvinium. LGK974 is a porcupine (PORCN) inhibitor. PORCN is a membrane-bound O-acyltransferase that is involved in the processing of Wnt ligand secretion. LGK974 was shown to inhibits Wnt signaling *in vitro* and *in vivo* (Liu et al., 2013). Pyrvinium (Vipyrvinium), is an FDA-approved anthelmintic drug used to treat pinworm. Pyrvinium is a potent inhibitor of Wnt signaling that inhibits Wnt signaling by binding to and increasing the activity of casein kinase 1α (CK1α) (Thorne et al., 2010).

To confirm that the selected drugs were exerting their effects on Wnt-signaling pathway in the sciatic nerve, we used the Wnt-signaling reporter line TCF/Lef:H2B-GFP (Ferrer-Vaquer et al., 2010). In this reporter line, six copies of the responsive element of the Wnt-signaling transcription factor Lef1 were inserted upstream of a gene that encodes a GFP-fused histone 2B protein. Active Wnt signaling pathway in this line results in GFP signal in the nucleus (Ferrer-Vaquer et al., 2010). Using this reporter line we found that crush injury induces Wnt signaling pathway activity in the sciatic nerve seven days post-injury. We found that as expected, treatment of mice with the Wnt signaling inhibitor LGK974 reduced Wnt-signaling activity (Figure 5A).

**Figure 5:**
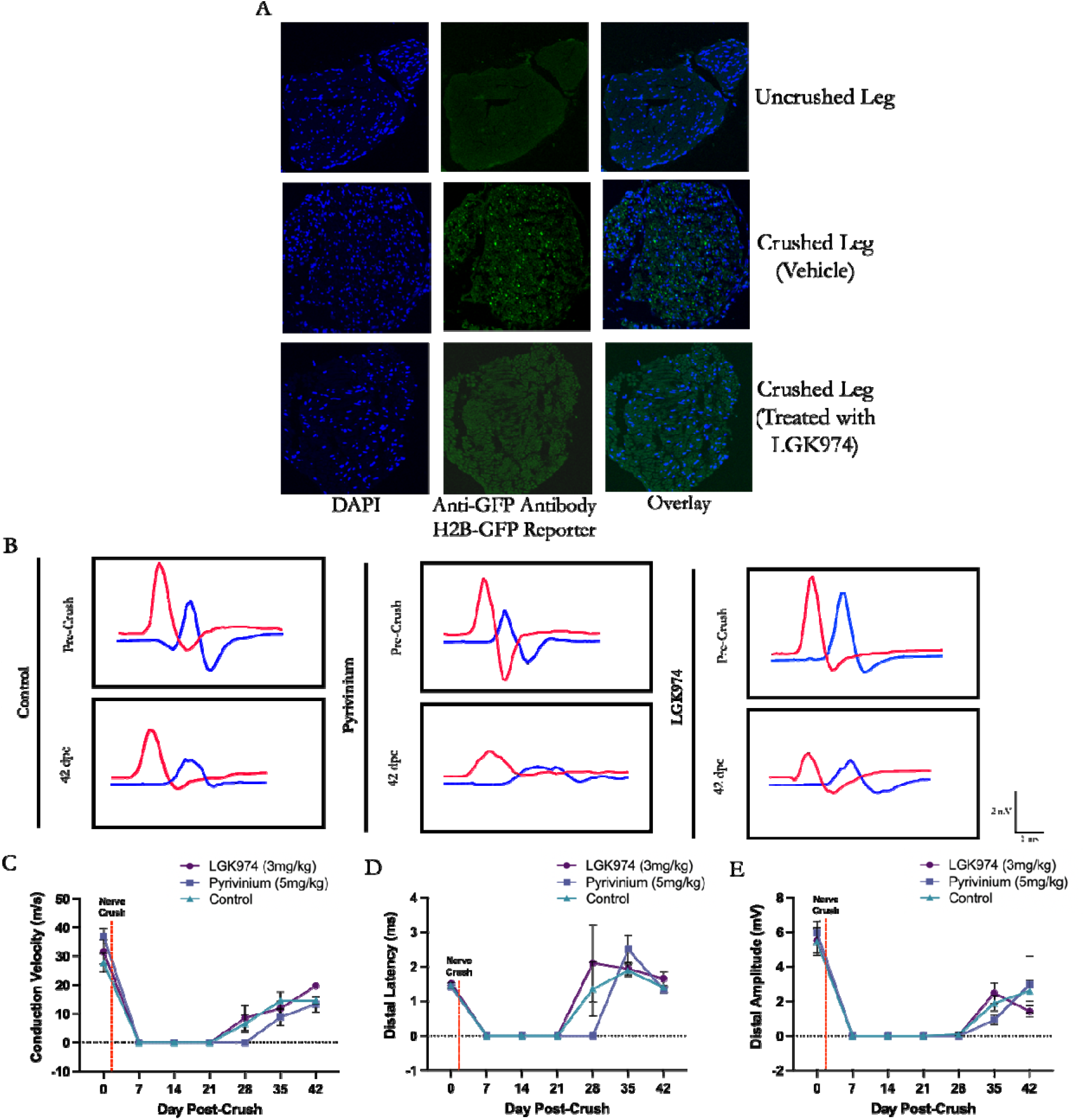
Pharmacological inhibition of the Wnt Signaling Pathway does not change sciatic nerve regeneration. Eight-week-old TCF/Lef:H2B-GFP mice were subjected to a nerve-crush injury. Immediately following crush surgery, mice were treated with Pyrvinium (5 mg/kg), LGK974 (3mg/kg), or vehicle for 7 days. Recovery was monitored through electrophysiological measurements of sciatic nerve compound motor action potentials (CMAPs). **(A)** The expression of the H2B-GFP reporter protein was only detected in the crushed leg, using an anti-GFP antibody (second row). Treatment of the TCF/Lef:H2B-GFP mice with LGK974 abolished expression of the reporter gene (third row). (B) Representative traces of electrophysiological findings in mice treated with Pyrvinium, LGK974, or vehicle at two time points - pre-crush and 42 days post-crush injury. Distal CMAP trace is in red, proximal CMAP trace is in blue. **(C-E)** Conduction velocity (m/s), distal latency (ms), and distal amplitude (mV) measurements through sciatic nerve for LGK974 (purple), Pyrvinium (blue), and vehicle-treated or (green) mice. There is demonstrated partial recovery of the sciatic nerve at 42 days post-crush for all mice, however, the metrics of functional recovery are not significantly different between mice in the three treatment groups. Note: In 5C, no error bar is displayed for LGK974 treatment at 42 days post-crush because the size of the error bar is smaller than the size of the symbol. Two-way ANOVA for multiple comparisons was used in the data analysis. Significance levels: P > 0.05 (n.s.), P ≤ 0.05 (*), P≤ 0.01 (**). N = 18 mice, 6 mice per treatment group.

To explore the effect of pharmacological inhibition of the Wnt Signaling pathway on the functional recovery of the sciatic nerve from the crush injury, we followed the functional recovery of the mice using EMG. We found that even though the Wnt signaling pathway is activated following crush-injury, inhibition of this pathway by either LGK974 or Pyrvinium did not change the functional recovery of sciatic nerve, as measured by electrophysiological examination of the sciatic nerve (Figure 5B-E). While there is variation between values obtained for distal amplitude (mV), distal latency (mS) and conduction velocity (m/s) at specific time points, treatment with Pyrvinium and LGK974 did not expedite or change the end-point recovery metrics when compared to the control (vehicle-treated), suggesting that Wnt inhibition with these drugs is not effective in improving recovery from nerve injury (Figure 5B-E).

## Discussion

In humans, PNS injuries fail to fully recover due to the inability of the axons to extend over long distances, resulting in denervation and atrophy of the distal tissue (Scheib and Höke, 2013). Therefore, finding new therapeutic targets that can expedite PNS regeneration is crucial. We and others have demonstrated that the Wnt signaling pathway controls developmental PNS myelination (Elbaz et al., 2016; Grigoryan et al., 2013). Recent studies suggest that Wnt signaling pathway is also re-activated upon PNS injury (Duraikannu et al., 2018; Vliet et al., 2021). Since the Wnt signaling pathway can be targeted by several drugs, we sought to investigate the therapeutic potential of Wnt signaling pathway to improve regeneration. In the present study, we used a combination of pharmacological and genetic approaches to manipulate Wnt signaling pathway during PNS regeneration. We found that neither genetic activation nor pharmacological inhibition of Wnt signaling pathway were able to expedite PNS regeneration. Our data therefore suggests that the Wnt signaling pathway is not a promising therapeutic target for enhancement of PNS regeneration.

To circumvent the effect Schwann-cell specific ablation of *Apc* on developmental PNS myelination, which we described previously (Elbaz et al., 2016), we used the inducible *PLP-CreER^t^* line (Doerflinger et al., 2003). To confirm recombination in Schwann cells, we used a reporter line (Figure 1). In agreement with previous studies that tested this line (Jessen and Mirsky, 2019, 2016), we found that the *PLP-CreER^t^* line mediates recombination mainly in non-myelinating Schwann cells. Nevertheless, recent single-nuclei RNA sequencing studies identified a sub-population of myelinating Schwann cells that express high levels of PLP (Yim et al., 2022). In line with these studies, we also found recombination in some myelinating Schwann cells. Further studies are required to determine the possible effect of *Apc* ablation in the *PLP-CreEP^t^* - *APC*^Lox/Lox^ mice on satellite glial cells, that also express PLP (Avraham et al., 2020). In addition, while we have focused here on the effect of Wnt signaling pathway manipulation on the motor performance and on the conduction velocity in the sciatic nerve, which reflects mainly the motor units, further studies are required to determine the possible effect of Wnt signaling pathway manipulation on the sensory neurons in the DRG, which are also are sensitive to the Wnt signaling pathway (Duraikannu et al., 2018)

Lineage tracing studies demonstrated that Schwann cells that are recombined in the *PLP-CreER^t^* line which was used here, actively participate in nerve repair (Jessen & Mirsky, 2019, 2016). Our genetic studies found that activation of the Wnt signaling pathway in this Schwann cell population by *Apc* ablation did not change the functional recovery of the PNS, suggesting that the PLP positive Schwann cells are not sensitive to Wnt signaling activation. The existence of a population of Mpz positive, PLP negative Schwann cells (Yim et al., 2022) may suggest that these Schwann cells, that were not recombined in our studies, may be sensitive to Wnt signaling manipulation. Nevertheless, the fact that the pharmacological treatments (that are not cell type specific) did not affect the PNS regeneration, argues against this and suggest that altering Wnt signaling does not change the functional recovery of the PNS.

In the CNS, remyelination is performed mainly by newly formed oligodendrocytes, derived from brain resident oligodendrocyte progenitor cells (OPCs). The differentiation of these cells upon remyelination is, in many ways, analogous to their differentiation during developmental myelination. Moreover, many of the transcriptional pathways that are important for developmental CNS myelination, are also implicated in CNS remyelination processes (Sock & Wegner, 2019). In sharp contrast, in the PNS, remyelination relies on the ability of mature Schwann cells to reprogram and turn into repair Schwann cells. These cells do not participate in developmental PNS myelination (Jessen & Mirsky, 2016, 2019). Hence, in the PNS regeneration is controlled primary by injury-induced transcriptional pathways that are different from the transcriptional pathways that controls developmental PNS myelination (Sock & Wegner, 2019). In line with this, while the Wnt signaling pathway is implicated in both CNS myelination and remyelination (Fancy et al., 2009), our study suggests that Wnt signaling pathway does not play a fundamental role in PNS remyelination, unlike in developmental PNS myelination in which Wnt signaling plays a fundamental role (Elbaz et al., 2016; Grigoryan et al., 2013). Although our studies suggest that Wnt signaling is not a strong therapeutic target for enhancement of PNS remyelination, further studies are required to determine the role of Wnt signaling pathway in Schwann-cell-mediated remyelination of the CNS. Upon CNS injury, Schwann cells derived from oligodendrocyte progenitor cells (OPCs) participate in remyelination of the CNS (Assinck et al., 2017; Crawford et al., 2016; Zawadzka et al., 2010). While the mechanisms that control OPC to Schwann cell differentiation are not understood, the ability of OPCs to generate Schwann cells upon CNS injury was attributed to local increase in Wnt and BMP4 signaling (Ulanska-Poutanen et al., 2018). This may suggest that the Wnt signaling pathway may play a role in Schwann-cell-mediated remyelination of the CNS.

The Wnt-signaling pathway is highly activated in human cancers, which has led to focusing on Wnt signaling pathway as a therapeutic target, and to the development of various Wnt signaling inhibitors (Jung & Park, 2020). PNS injuries lead to activation of Wnt signaling pathway in both human and rodents (Duraikannu et al., 2018; Vliet et al., 2021). Nevertheless, our data suggest that Wnt signaling pathway is not a good therapeutic target for enhancement of PNS regeneration.

## Author contributions

N.M. designed research, performed research, analyzed data and wrote the paper; M.V: performed research. B.E.: designed research, performed research, analyzed data, wrote the paper.

## Acknowledgments

BE is supported by the NINDS (R01 NS067550), by The National Multiple Sclerosis Society (RG-1501-02797) and by the Department of Defense Office of Congressionally Directed Medical Research (W81XWH2210386).

